# Universal allometry from empirical parameters

**DOI:** 10.1101/2021.05.20.444891

**Authors:** Jody C. McKerral, Maria Kleshnina, Louise Bartle, James G. Mitchell, Jerzy A. Filar

## Abstract

Allometric settings of population dynamics models are appealing due to their parsimonious nature and broad utility when studying system level effects. Here, we parameterise the size-scaled Rosenzweig-Macarthur ODEs to eliminate prey-mass dependency. We define the functional response term to match experiments, and examine situations where metabolic theory derivations and observation diverge. We produce dynamics consistent with observation. Our parameterisation of the Rosenzweig-Macarthur system is an accurate minimal model across 15+ orders of mass magnitude.

## 1. INTRODUCTION

Allometric scaling relationships have been the subject of scrutiny and debate since the connection between organism size and its metabolic rate was first defined by Rubner in 1883 [1–4]. These models, which link some characteristic *y* to the size *x* of an organism via the power law *y* = *ax^b^* (where *a*, *b* are scalar constants), are appealing due to their capacity to capture a multitude of relationships despite their simplicity. Scaling laws have been used to express a variety of biological rate measures, such as metabolism, consumption, and birth or death rates [5–8]. Allometry is also utilised in modelling behavioural traits and bioenergetic characteristics, such as movement behaviour or locomotory costs [7, 9–11]. At broad scales, such laws have been applied to ecosystem-level properties, including predictions of organism population density and carrying capacity [12–14]. However, despite scaling laws’ wide utility and intensive study, there has been a limited exploration of the properties of minimally constructed, size-generalised predator-prey models [15–17].

Many authors have examined the empirical relationship between organism and population sizes [12, 13, 18–20]. Reported exponents fall between −1 and −1/4 depending on factors such as taxonomy or environment. The classical −3/4 value describing global size-density relationships is the direct inverse of Kleiber’s 3/4 law for metabolic scaling [2, 13], leading to the ‘energetic equivalence’ hypothesis: that is, the net energy contained within each size class is invariant [18, 20]. This conjecture has been widely debated, particularly with respect to whether this invariance is cause or effect of other bioenergetic drivers [21]. However, despite disagreement over underlying mechanisms, there is broad consensus that the consistency of size-density scaling within empirical likely reflects fundamental physical constraints [5, 21]. To examine what drives limitations in macro-scaling behaviour, it is possible to use dynamical size-based models incorporating organism traits that scale across the size range [6, 15, 16]. This approach facilitates the investigation of critical breaks in ecosystem-level scaling laws within a global framework, and the exploration of potential impacts from changes that may affect many organisms in a similar way – for example, warming temperatures or emergent hypoxia in the oceans [22, 23]. However, perturbing parameters across 15+ orders of magnitude in size poses challenges. For example, coexistence regions of size-generalised predator-prey models are dominated by scaling exponents [16].

It is more straightforward to keep model behaviour stable, thus resolving the coexistence issue, by using the 4-parameter Lokta-Volterra model, but that setting is too simple for some applications [6, 17]. Alternately, a series of models may be solved piece-wise for different size classes, yet this means that they are not truly generalised. In the most comprehensive study to date, a size-based paramaterisation of the Rosenzweig-Macarthur system places restrictions on the relationships between the exponents of each parameter [16]. However, this in turn limits the types of perturbations that may be applied or investigated. Finally, there are discrepancies in the treatment of the functional response term between the theoretical and empirical literature. The theoretical literature broadly assumes that the limit of maximal consumption ties predator production to the prey’s and thus scales negatively to match prey production, however, there is empirical biological evidence for positive scaling.

Here, we present an alternate approach of paramaterising size-based predator-prey interactions for the classical Rosenzweig-Macarthur differential equations. Under this framework, we examine the parameter sensitivity and required ranges for species coexistence within the context of real-world observations. We are able to show that, despite the number of assumptions inherent within this style of modelling, the mathematical restrictions are closely related to biological observations. Finally, we describe the conditions required to create an entirely size-invariant model, and how empirically derived parameters generate ODE solutions that match real-world size-abundance distributions.

## 2. PARAMETERISATION OF THE MODEL

We begin with the Rosenzweig-Macarthur ODEs [24] and Holling II functional response,

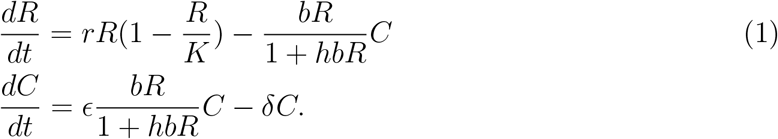

We have variables *R* for resources and *C* for consumers. The parameters *r* and *δ* are birth and death rates respectively. Carrying capacity is given by *K*, interaction rate *b*, handling time *h* and the conversion efficiency *ϵ*. To investigate the system across the full size range we scale the parameters by mass. Organism size (in g) is given by *S_R_* for resources and *S_C_* for consumers. We depart from [16] by constructing the functional response term in line with parameterisations used within experimental research [6, 25, 26]. Hence, the strictly positive parameters are expressed as

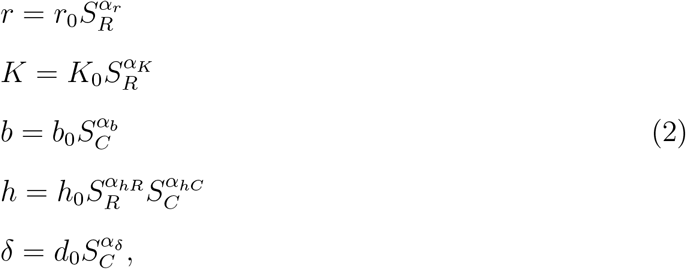

where for each parameter *i*, the coefficients *i*_0_ may be standardised to a boundary value, and *α_i_* denotes the scaling exponent. Next, we define the prey-predator mass ratio as *ρ*, where *ρ* > 0. We may then relabel the parameters *r*, *h* and *K* in terms of the consumer,

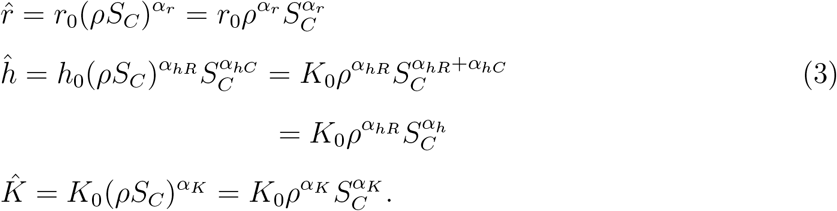

With this approach we extend the results of [16] by placing no restrictions on the exponents, allowing *h* to be an independent term which may be matched to empirical observations. We now also follow standard practice by setting *ϵ* ∝ *ρ*, that is, the conversion efficiency is proportional to the prey-predator mass ratio [16]. Next, we use a standard rescaling of (1) to reduce the number of parameters and simplify analyses. We set 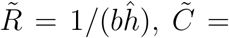 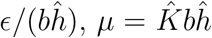, and define 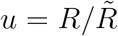 and 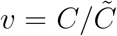. After scaling time by 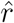, and defining 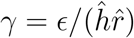 and 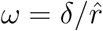, we arrive at the new system

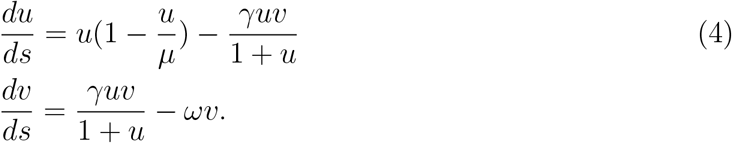

The parameters in (4) are also all strictly positive and scale across the size range. For completeness, we provide the explicit relationship between the old and new parameters in Table I, and the system of equations with substituted terms is

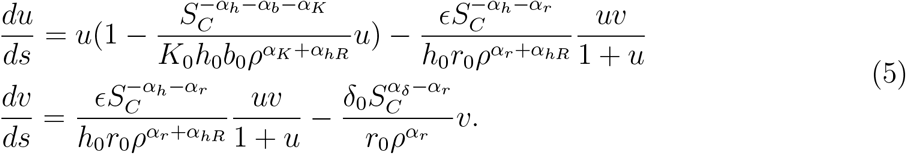

**TABLE I.** Relationship between parameters in original and rescaled Rosenzweig-Macarthur system. Here, *α_h_* = *α_hR_* + *α_hC_*.

|  | Definition | Coefficient | Exponent |
| --- | --- | --- | --- |
| $\mu$ | $\hat{K}\hat{h}b$ | $\mu_0 = K_0 h_0 b_0 \rho^{\alpha_K + \alpha_{hR}}$ | $\alpha_\mu = \alpha_K + \alpha_h + \alpha_b$ |
| $\gamma$ | $\epsilon/\hat{h}\hat{r}$ | $\gamma_0 = \epsilon/(r_0 h_0 \rho^{\alpha_r + \alpha_{hR}})$ | $\alpha_\gamma = -\alpha_h - \alpha_r$ |
| $\omega$ | $\delta/\hat{r}$ | $\omega_0 = \delta_0/(r_0 \rho^{\alpha_r})$ | $\alpha_\omega = \alpha_\delta - \alpha_r$ |

The expression *S_R_* = *ρS_C_* facilitates interpretability in downstream analyses. All exponent terms may be collected within *S_C_*, which we henceforth refer to as *S*. This provides simplified expressions within Table I and (5), yet we may still examine the impacts of perturbations to any one parameter. Table II summarises parameter exponent ranges within empirical research.

**TABLE II.** Literature bounds on parameter values. The top portion of the table outlines scalars. The middle section summarises scaling exponents, and the bottom the exponent range for (4). Value limits are given for completeness. However, there is consensus that *α_r_, α_d_* ≃ −1/4, also verified in a substantial recent review [27]. Similarly, despite the potential range for *α_b_*, typically 1/2 ≤ *α_b_* ≤ 1, significantly constraining the exponent ranges in (4).

| Symbol | Parameter | Minimum | Maximum | References |
| --- | --- | --- | --- | --- |
| $\rho$ | Prey-predator mass ratio | 1E-4 | 1E2 | |
| $S$ | (Consumer) mass, g | 1E-10 | 1E7 | |
| $\epsilon$ | Conversion efficiency | 0 | $\rho$ | |
| $\alpha_r$ | Birth rate | -0.81 | -0.25 | [6, 15, 16, 22, 27–31] |
| $\alpha_\delta$ | Death rate | -0.35 | -0.22 | [6, 15, 16, 22, 27–31] |
| $\alpha_b$ | Interaction rate | -0.25 | 1.58 | [6, 17, 25, 26, 30] |
| $\alpha_K$ | Carrying capacity | -0.88 | -0.74 | [6, 16, 31] |
| $\alpha_{hR}$ | Handling time (resource) | 0 | 1 | [15, 25, 26] |
| $\alpha_{hC}$ | Handling time (consumer) | -1.1 | 0 | [6, 15, 25, 26] |
| $\alpha_\mu$ | | -2.2 | 1.84 | |
| $\alpha_\gamma$ | | -0.75 | 1.92 | |
| $\alpha_\omega$ | | -0.1 | 0.59 | |

## 3. RESULTS

### 3.1. Coexistence & Sensitivity

The non-trivial equilibrium of interest (coexistence) is obtained by equating the right side of (4) to zero and solving for *u* = *u*\* and *υ* = *υ*\*, yielding

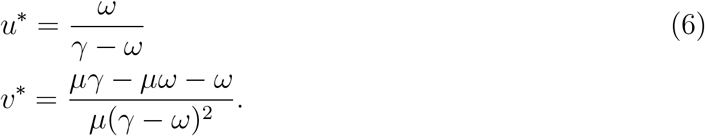

For there to be non-negative values for (*u*\*, *υ*\*), we require that

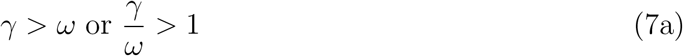

and

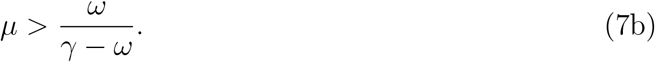

For the Jacobian of the right side of (4), the condition *det* > 0 is also fulfilled by (7b). We may express (7b) as *γ/ω* > 1+1/*μ*. As all parameters are strictly positive, if (7b) is satisfied, it immediately follows that (7a) is satisfied also. The inequality

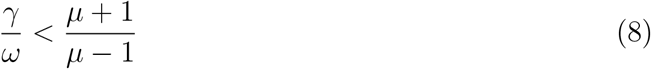

determines the sign of the trace of the aforementioned Jacobian, which dictates whether the system converges to a point or to a stable limit cycle, and there is a Hopf bifurcation at equality. The dynamical characteristics of the Rosenzweig-Macarthur system have been explored in depth elsewhere (such as [24, 32, 33] and references within). Our focus is the interplay between biological and mathematical constraints. We now discuss inequalities (7)–(8) in the context of empirical observations.

#### 3.1.1. Handling time

The condition from (7a) is equivalent to

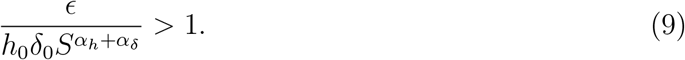

Examining the exponents, if *α_δ_* + *α_h_* < 0, the condition may fail for small organisms. As *α_δ_* is tightly constrained (Table II), we examine system behaviour when varying the scaling of handling time.

Handling time’s classical null model from Yodzis & Innes [15] based on metabolic theory is equivalent to 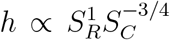, or 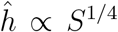 (as derived in [25]^1^). It results in a maximal consumption rate of *c* ∝ *S*^−1/4^. This matches the setting of [16], where the birth rate of the predator in the presence of unlimited resources is assumed to scale with the birth rate of prey. However, consideration of physiological traits’ nuanced impact on handling time has since led to a departure from the traditional metabolic framework. Attacking, killing, then eating and digesting prey all impact the parameter *h* [26] and the prey’s contribution to the process should be incorporated [25, 26]. Handling time has therefore been recast to the more biologically representative form we use in this study: 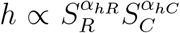, where typically 0 ≤ *α_hR_* ≤ 1 and −1 ≤ *α_hC_* ≤ 0 [25, 26]. The exponents *α_hR_*, *α_hC_* have been empirically determined in several reviews and display considerable variability [6, 25, 26]. We now discuss the implications this variability has for coexistence under the inequalities in (7a)–(7b).

Two of the three reviews listed above conclude that the predator-prey components of handling time scale more gently (whether positive, or negative) than null models predict. However, the resultant exponent for 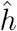 is positive (≃ 1/3) for arthropod functional responses [25], but negative (≃ −1/8) when examining broader taxonomic groups [26]. Most of the organisms in [26] display negative scaling for 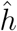 in taxa-specific breakdowns due to gentler scaling of the resource exponent. Only *α_hC_* is assessed in [6], which is calculated across a wide range of taxa for 2D and 3D environments and for a larger mass range than in [26]; for the 2D and 3D case *α_hC_* ≃ −1.1. The authors account for the steeper scaling relative to metabolic expectation by noting that feeding is an active process scaling with maximal rather than basal metabolism [6]. To calculate the scaling of 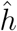 from the empirical assessment in [6], we use the assumption *α_hR_* = 1, which is the parameter’s upper limit. This implies the exponent *α_h_* ≤ −0.1, and that 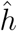 scales below the value of 1/4 assumed by previous theoretical work on the model. Conceptually, this indicates that the parameter may be constrained by physical processes rather than a bioenergetic flux balance. Note that despite the phenomenological formulation of the functional response predator production is implicitly constrained by the prey density. Coexistence condition (7b) may be expressed as *γ/ω* > 1 + 1/*μ* meaning ln(*γ/ω*) > ln(1 + *μ*^−1^); if we then substitute the original parameters we arrive at the inequality

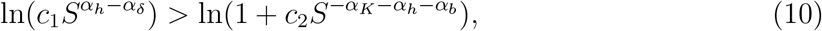

where *c*_1_, *c*_2_ are constants derived from the coefficients. Considering (10) together with the empirical behaviour of 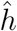, if *α_h_* < −1/5 across the size range, smaller organisms may violate this condition (Figure 2). If *α_h_* ≃ −*α_δ_* then all sizes will fulfil the condition. There is empirical support for these observations. The taxonomic group breakdowns in [26] indicate that smaller taxa may display positive scaling for 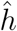 as concluded in [25] and the strongly negative scaling is observed in macro-organisms, particularly vertebrates. The notable exception of unicellular marine organisms (*α_h_* ≃ −1/3) has a very small sample size; further experimental study may therefore reach an alternate conclusion.

#### 3.1.2. Carrying capacity and scaling of population cycling

As the right side of (7b) is equal to *u*\*, the coexistence condition may also be expressed as

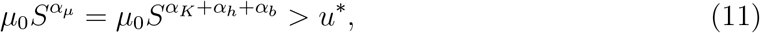

where 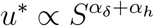. If we combine (7b) and (8), we obtain

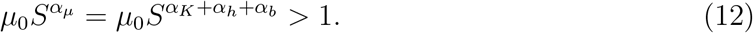

Together, (11) and (12) indicate the carrying capacity must be sufficiently high for a sustainable prey population, and that predator attack rates must be high enough to compensate for mortality across all sizes. Assuming reasonable values for *α_K_*, *α_b_* such as those given in Table II, these inequalities will generally hold. Under the original system (1) and assuming coexistence, resource equilibria will scale with size as 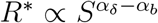 and consumer equilibria will scale as 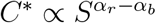. Despite their importance for the coexistence domain, the carrying capacity and half saturation do not meaningfully impact equilibria abundances in allometric paramaterisations. However, they do impact some properties of the limit cycle.

Allometric settings of the Rosenzweig-Macarthur system usually result in oscillating solutions due to size-scaled parameter values relative to the constraints in (8) [29]. We find stronger empirical support than in previous work for a size scaling signal for the period *τ* [29]. The theoretical scaling 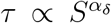 (derived in both [16, 29], with a similar result in [15]) agrees well with observed values (Figure 1b). A previous review has found that the ratio of maximum to minimum densities is size-invariant [29], indicating that the oscillation amplitude decreases with increasing size. Further qualitative support that this mathematical behaviour is aligned with features of real-world biological systems is given by the fact that log-transformed size-density relationships demonstrate near-constant variance across 15+ orders of magnitude [27, 43]. Under our paramaterisation the log-scaled oscillations are relatively sinusoidal and may be constrained between one to three orders of magnitude (Figure 1a). Hence, they are more realistic than allometric Lokta-Volterra dynamics which push predator populations to unreasonably low levels with fluctuations exceeding 15 orders of magnitude [17]. Unfortunately early efforts to find analytic approximations of the oscillation amplitude of the Rosenzweig-Macarthur system have not been generalised [44]. Recent results have only been derived for specific – and restricted – parameter values [45]. However, the rescaled system (4) provides scope for us to examine the effects of perturbations in a simplified manner. In Figure 1c, we show through simulation that perturbations to all parameters may impact the magnitude of the fluctuation of the limit cycle. However, unless these perturbations are applied to *α_r_* or *α_δ_*, the oscillation amplitude will remain (nearly) invariant with respect to the mean population density. We next use a sensitivity analysis to assess the system’s robustness to different forms of perturbation in Section 3.1.3.

**FIG. 1.**
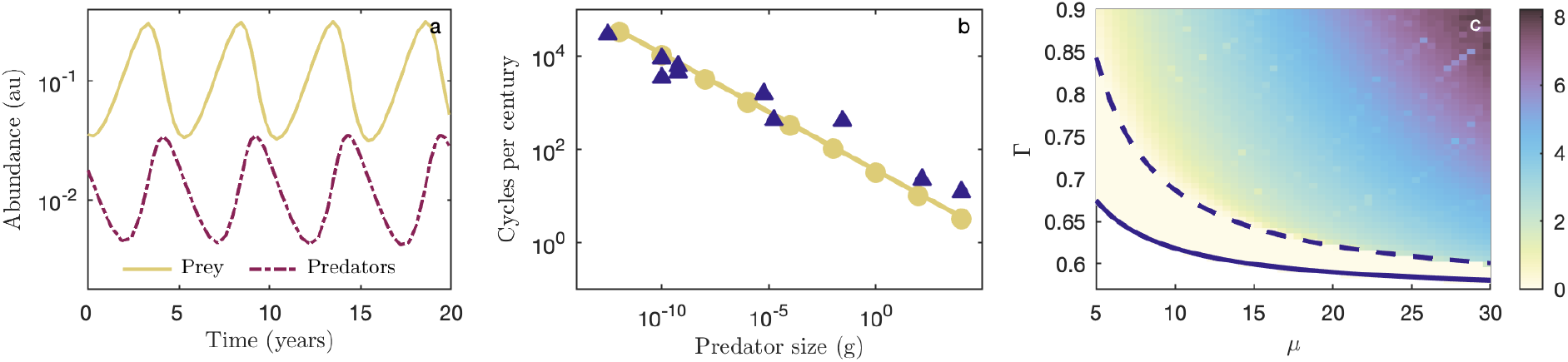
Properties of the limit cycle: (a) Predator (purple)-prey (yellow) oscillations for a 10g predator (b) Limit cycle period scaling for numerical (circles), empirical (triangles) and analytic (solid line) results. Only predators are shown. Data is from population study time series [17, 34–41]; periods were calculated by using supplied data files or extracting data from figures [42], and averaging time between peaks. (a)-(b) use empirical scaling of 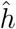, where *α_h_* = −1/8. (c) Dynamics of the rescaled system under different parameter values. Here, Γ = *γ/ω*, which is plotted against *μ*. The region below the solid line indicates no coexistence, between the solid and dashed lines denotes a sink to the equilibrium point, and above the dashed line a stable limit cycle. Colour indicates the difference between the (log) maximum and minimum predator abundances.

**FIG. 2.**
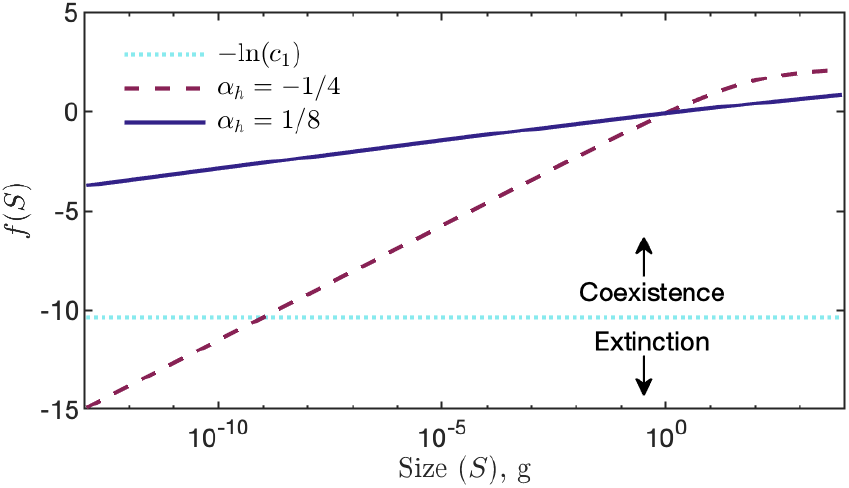
Graphical representation of coexistence condition (7b). Rearranging (10): 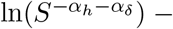 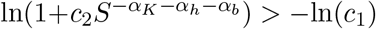. We denote the left side of the inequality as *f* (*S*); if *f* (*S*) > −*ln*(*c*_1_), there is coexistence. Under the feasible values in Table II, here *α_K_* and *α_b_* have less effect than *α_h_*; we thus set *α_K_* = −3/4 and *α_b_* = 1/2.

#### 3.1.3. Sensitivity

A local sensitivity analysis provides a first-order approximation of the relative impact of changing parameters on the solutions of (4) near the system’s equilibria. We adhered to the methodology described in [46]. In order to check how sensitive the system, 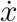, is to small changes in parameters, *λ_i_*, we construct a sensitivity function, *S*(*t*), such that

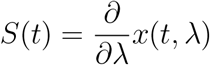

and *x*(*t, λ*) is a solution of 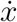. Next, we characterise the solution to the sensitivity equation given by

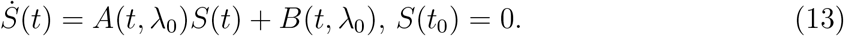

Applying (13) to (4), *A* is the Jacobian of (4) with respect to variables *u* and *υ*, and *B* is the Jacobian of (4) with respect to parameters *μ*, *γ*, and *ω*, both of which are evaluated at nominal parameter values. After setting initial conditions *u*_0_ and *υ*_0_, we obtain numerical solutions for (13). Figure 3 shows the trajectories for two initial conditions. Firstly, for *u*_0_ and *υ*_0_ in the neighborhood of *u*\*, *υ*\* respectively, and secondly for *u*_0_, *υ*_0_ an order of magnitude greater/smaller than *u*\*, *υ*\* respectively. The qualitative behaviour remains the same in both cases. For an initial state near the equilibrium (Figure 3(a-b)), there is monotonic behaviour as the system converges to the limit cycle. In the case of Figure 3(c-d) the limit cycle emerges after some critical time *t_c_*. For each, the system is least sensitive to *μ*, providing further support that perturbations to *r* and *δ* have the largest impacts on the system. The qualitative behaviour is similar for other nominal parameter values, provided they are not set on the other side of the bifurcation boundary. The exponents with the least empirical variation – by a significant margin – are *α_r_* and *α_δ_*, mirroring the mathematical constraints. That is, the dynamical behaviour of the system is relatively robust to perturbing the functional response parameters displaying the highest empirical variance.

**FIG. 3.**
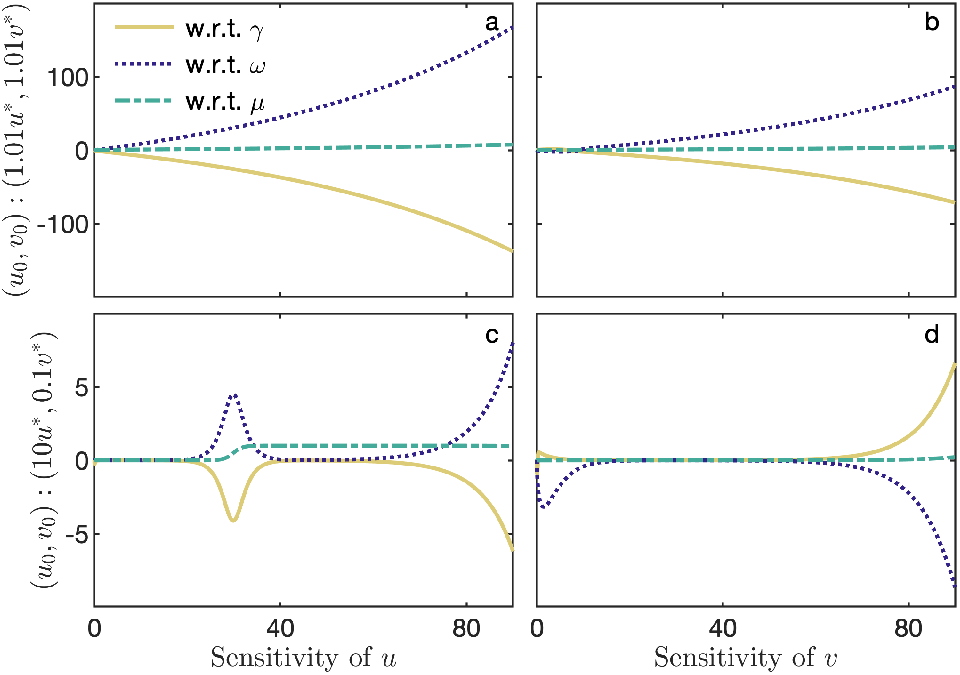
Sensitivity of rescaled system. *x*-axis denotes time (au). (a-b): Sensitivity of the solutions of (5) to perturbations to each of the parameters under an initial condition near the point (*u*\*, *υ*\*). (c-d): As above, except under an initial condition (10*u*\*, 0.1*υ*\*); the trajectory also converges to the limit cycle.

### 3.2. Applications

The rescaled parameter definitions in Table I may be used to determine a size-invariant system by balancing the exponents. This is desirable as it is straightforward to set coexistence for an arbitrary size range and facilitates analytic study of the equations. By the definition of 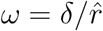, the size scaling of *r* must match *δ*, both of which consistently display an exponent of −1/4. It immediately follows that *α_h_* = 1/4 as 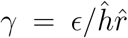. Assuming that carrying capacity scales with a −3/4 exponent, and using the definition 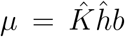 it follows that *α_b_* = 1/2. As the dynamics of (4) are identical to (1), eliminating the size-dependency in (4) is mathematically expedient and we note that the majority of allometric studies to date have used this approach. However, given the tenuous empirical support for 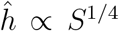, it may be more biologically sound to assign a value where *α_h_* ≤ 0. A full size-abundance distribution may still be generated when parameter scaling is set to empirical values. A recent size-density scaling analysis indicates that the relationship follows *N* * ∝ *S*^−1^ rather than the canonical *N* * ∝ *S*^−3/4^ (where *N* * is population density). Indeed, using empirical scaling of *b* results in a distribution close to *N* * ∝ *S*^−1^ (Figure 5). Whilst the lower bound of *α_b_* ≃ 1/2, estimates of the ‘universal’ value suggest 0.6 < *α_b_* < 0.9. A limitation of our treatment of *b* is that we place no restrictions on the interactions between a predator and any arbitrary-sized prey. Predator-prey interaction processes are complex and increasing evidence suggests they follow a ‘hump-shaped’ curve with the predator-prey mass ratio [25]. A natural extension to our model would be to introduce a term reliant on *ρ* to the parameter *b*, where 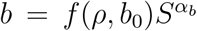, and *f* (*ρ, b*_0_) is a function assigning probability of prey capture based on the prey-predator size ratio. While a form for *f* (*ρ*) has been proposed [16], it would be possible to use a function encoding a broader range of life history traits for the tradeoff of introducing more parameters. Relevant processes to consider may include habitat effects on foraging, prey refuges, and optimal size ratios, resulting in further constraints on coexistence.

Our final consideration is the contribution of *ρ* to the conversion efficiency *ϵ*. The empirical distribution of *ρ* is approximately lognormal peaking at ≃ 0.02 [30]. Equilibria population ratios do not follow a 1:1 relationship with the size ratio of prey and predator when using the relationship *ϵ* = *ρ* (solid white line, Figure 4). For a fixed predator size and increasing prey size, organisms become less efficient at converting biomass. However, this result does not align with observed data. A review of 15,000+ predator-prey pairs concludes that size differences between predator and prey has an upper limit, potentially due to inefficiencies when the size discrepancy becomes too extreme [48]. Furthermore, the larger the predator, the more generalist its feeding strategies [25]; increases in prey biomass – which could indicate predators feeding on smaller prey – do not translate to a proportionate increase in predator biomass [47]. We therefore apply the assumption that energetic reward (and biomass conversion) for predator effort declines as the size difference increases, and that the scaling of *ρ* with equilibria population ratios is superlinear. We can implicitly capture the result by assigning a function *ϵ* = *aρ^ψ^*, where *ψ* is a scalar. For simplicity, we set *a* to 1, and in Figure 4, we assess the predator-prey population ratios for varying *ψ*. A value of *ψ* > 1 increases the difference between the predator-prey populations; *ψ* < 1 reduces it. More sophisticated functional forms may include favourable size ratios or introduce a size dependency to the value of *ϵ*. However, there is limited empirical research on scaling properties of *ϵ* [13, 19, 28]. A theoretical investigation of optimal predator-prey size ratios together with more complex functional response formulations reflecting alternate foraging/feeding strategies may yield interesting results. We leave this question open for future work.

**FIG. 4.**
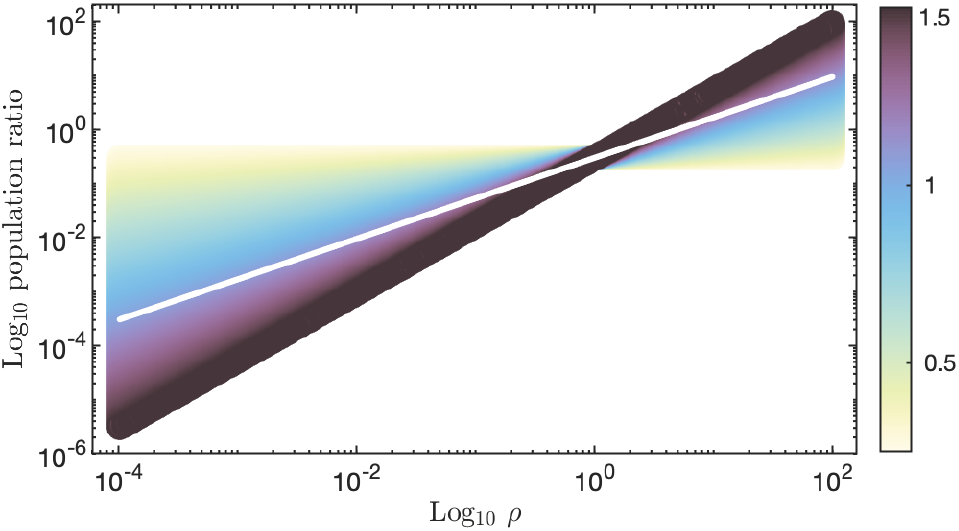
Impact of perturbing conversion efficiency *ϵ* by setting a function on *ρ*. Here, *ϵ* = *ρ^ψ^*, where −1/2 < *ψ* < 3/2. Colours map to the value of *ψ*, and the white line depicts *ψ* = 1. Y-axis shows the ratio of the predator-prey equilibria populations. Result is size invariant.

To generate the size-abundance distribution shown in Figure 5, we use a value of *ψ* ≃ 1.3, together with scaling values of *α_r_* = *α_δ_* = −1/4, *α_h_* = −0.1, *α_K_* = −3/4 and *α_b_* = 2/3. The model’s distribution scales to −0.92, close to the −0.95 value observed in [27]. The inset shows the predator-prey density relationship, scaling at 0.76, close to the ≃ 3/4 findings in [47]. Note that to generate Damuth’s law, a value of *ψ* = 1.3 and *α_b_* = 1/2 results in exponents of −3/4 and 3/4 for the size-abundance and predator-prey density scaling respectively.

**FIG. 5.**
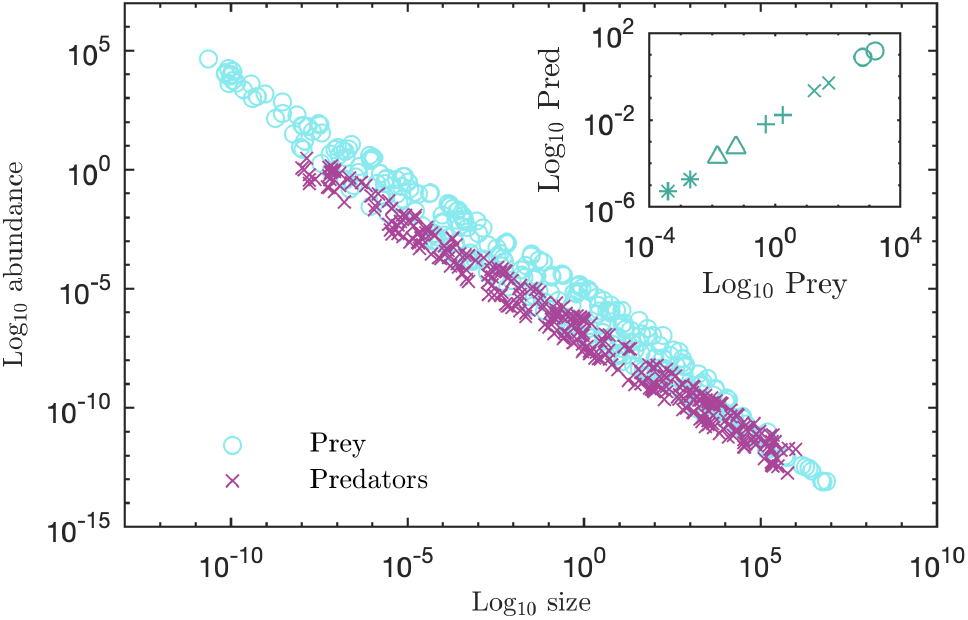
Main: Size-abundance data generated from the model. Circles depict prey abundances, and crosses predators. The parameter *ρ* was randomly selected within the interval of 1E-4 and 1E2. The full size-abundance distribution from the model scales to −0.91, close to the empirical distribution in [27] of −0.95. Inset: each pair of points are the maximum and minimum abundances attained by the predator/prey during limit cycle oscillations. We show examples of five predator sizes: 1E-6 (○), 1E-4(×), 1E-2 (+), 1E0 (Δ), and 1E2g (∗). For each *ρ* = 0.02. The slope within each symbol group ≃ 0.76. That is, increasing prey density does not result in a 1:1 increase in predator density. The scaling relationship is sublinear, matching the observations of [47].

## 4. CONCLUSIONS

Here, we investigate the links between empirical and theoretical allometric literature. The resource size dependency is eliminated from the system by explicitly encoding the prey-predator mass ratio, *ρ*. This simplifies analyses and provides a parsimonious base for customising the equations, which may be useful for food web or trophic modelling. We find that the model constraints complement empirical observation. Contrary to most previous studies, we use an empirically determined parameterisation of the functional response term. Our results suggest that the standard approach of setting exponents based on metabolic theory may need to be reassessed. The handling time parameter shows the greatest departure from those assumptions, and the highest variance, which is consistent with the massive trait variation in hunting and feeding strategies. Nevertheless, we find that results generated from an empirical setting agree with results in recent reviews of size-abundance scaling. This work may be extended in several ways. Firstly, one could incorporate temperature effects, for example after [22, 49], which may further stabilise the model by reducing the interaction strengths [28]. Secondly, additional empirical data on functional responses at the size extrema could more accurately define the scaling of 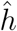 and *b*. Type I, Type III, or generalised functional responses may also be examined, although we note that system behaviour usually remains qualitatively similar over large size ranges [6, 31].

The broad limitation of allometry is that a generalist strategy can be a poor predictor of taxon-specific outcomes. Challenges to the framework arise not only from biological differences but also from physical or spatial processes, such as prey patchiness or heterogeneous habitat distribution [50]. Thus, care over interpretation and the applicability of results must be taken, particularly at the size limits in either direction. For example, prokaryotic reproduction rates fall between minutes and millenia [51, 52]. Furthermore, large organisms such as whales play a critical role in nutrient recycling; assuming a single species may be defined as a resource or consumer alone does not account for the intrinsic complexities within natural environments [53]. However, despite these caveats, allometric approaches have been found to outperform those explicitly encoding organisms’ individual and life-history traits when investigating a system’s macro properties [25]. Classical population dynamics models remain a powerful tool in ecology, and the consistency across many allometric laws suggest self-organising processes we are yet to unravel. We propose that systematically assessing where theoretical and empirical properties of allometric modelling diverge may assist in identifying plausible mechanisms governing these phenomena.

## Footnotes

1 The derivation assumes predator consumption (the inverse of handling time) scales with metabolic demand 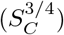, and the per-prey metabolic demand is therefore 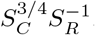. This matches the assumptions of [16, 17], however we note that other interpretations of the same model either do not normalise against prey mass (e.g. [31]) or do so implicitly (e.g. [6]).

## References

[1] M. Rubner, Ueber den einfluss der korpergrosse auf stoffund kaftwechsel, Zeitschrift fur Biologie 19, 535 (1883).

[2] M. Kleiber, Body size and metabolism, ENE 1, E9 (1932).

[3] G. B. West, J. H. Brown, and B. J. Enquist, The fourth dimension of life: fractal geometry and allometric scaling of organisms, Science 284, 1677 (1999).

[4] J. L. Blanchard, R. F. Heneghan, J. D. Everett, R. Trebilco, and A. J. Richardson, From bacteria to whales: Using functional size spectra to model marine ecosystems, Trends in Ecology & Evolution (2017).

[5] G. B. West and J. H. Brown, The origin of allometric scaling laws in biology from genomes to ecosystems: towards a quantitative unifying theory of biological structure and organization, Journal of experimental biology 208, 1575 (2005).

[6] S. Pawar, A. I. Dell, and V. M. Savage, Dimensionality of consumer search space drives trophic interaction strengths, Nature 486, 485 (2012).

[7] R. Bale, M. Hao, A. P. S. Bhalla, and N. A. Patankar, Energy efficiency and allometry of movement of swimming and flying animals, Proceedings of the National Academy of Sciences 111, 7517 (2014).

[8] M. J. Reiss, The allometry of reproduction: why larger species invest relatively less in their offspring, Journal of Theoretical Biology 113, 529 (1985).

[9] D. W. Sims, E. J. Southall, N. E. Humphries, G. C. Hays, C. J. Bradshaw, J. W. Pitchford, A. James, M. Z. Ahmed, A. S. Brierley, M. A. Hindell, et al., Scaling laws of marine predator search behaviour, Nature 451, 1098 (2008).

[10] J. P. Haskell, M. E. Ritchie, and H. Olff, Fractal geometry predicts varying body size scaling relationships for mammal and bird home ranges, Nature 418, 527 (2002).

[11] J. G. Mitchell, The energetics and scaling of search strategies in bacteria, The American Naturalist 160, 727 (2002).

[12] J. Damuth, Population density and body size in mammals, Nature 290, 699 (1981).

[13] E. P. White, S. M. Ernest, A. J. Kerkhoff, and B. J. Enquist, Relationships between body size and abundance in ecology, Trends in ecology & evolution 22, 323 (2007).

[14] J. R. Bernhardt, J. M. Sunday, and M. I. O’Connor, Metabolic theory and the temperature-size rule explain the temperature dependence of population carrying capacity, The American Naturalist 192, 687 (2018).

[15] P. Yodzis and S. Innes, Body size and consumer-resource dynamics, The American Naturalist 139, 1151 (1992).

[16] J. S. Weitz and S. A. Levin, Size and scaling of predator–prey dynamics, Ecology letters 9, 548 (2006).

[17] A. Eilersen and K. Sneppen, Applying allometric scaling to predator-prey systems, Physical Review E 99, 022405 (2019).

[18] A. P. Allen, J. H. Brown, and J. F. Gillooly, Global biodiversity, biochemical kinetics, and the energetic-equivalence rule, Science 297, 1545 (2002).

[19] J. P. DeLong and D. A. Vasseur, Size-density scaling in protists and the links between consumer-resource interaction parameters, Journal of Animal Ecology 81, 1193 (2012).

[20] J. Damuth, A macroevolutionary explanation for energy equivalence in the scaling of body size and population density, The American Naturalist 169, 621 (2007).

[21] N. J. Isaac, D. Storch, and C. Carbone, The paradox of energy equivalence, Global Ecology and Biogeography 22, 1 (2013).

[22] V. M. Savage, J. F. Gillooly, W. H. Woodruff, G. B. West, A. P. Allen, B. J. Enquist, and J. H. Brown, The predominance of quarter-power scaling in biology, Functional Ecology 18, 257 (2004).

[23] A. Yool, A. P. Martin, T. R. Anderson, B. J. Bett, D. O. Jones, and H. A. Ruhl, Big in the benthos: Future change of seafloor community biomass in a global, body size-resolved model, Global Change Biology (2017).

[24] M. L. Rosenzweig and R. H. MacArthur, Graphical representation and stability conditions of predator-prey interactions, The American Naturalist 97, 209 (1963).

[25] G. Kalinkat, F. D. Schneider, C. Digel, C. Guill, B. C. Rall, and U. Brose, Body masses, functional responses and predator-prey stability, Ecology letters 16, 1126 (2013).

[26] B. C. Rall, U. Brose, M. Hartvig, G. Kalinkat, F. Schwarzmüller, O. Vucic-Pestic, and O. L. Petchey, Universal temperature and body-mass scaling of feeding rates, Philosophical Transactions of the Royal Society B: Biological Sciences 367, 2923 (2012).

[27] I. A. Hatton, A. P. Dobson, D. Storch, E. D. Galbraith, and M. Loreau, Linking scaling laws across eukaryotes, Proceedings of the National Academy of Sciences 116, 21616 (2019).

[28] M. M. Osmond, M. A. Barbour, J. R. Bernhardt, M. W. Pennell, J. M. Sunday, and M. I. O’Connor, Warming-induced changes to body size stabilize consumer-resource dynamics, The American Naturalist 189, 718 (2017).

[29] A. J. Hendriks and C. Mulder, Delayed logistic and rosenzweig–macarthur models with allometric parameter setting estimate population cycles at lower trophic levels well, Ecological complexity 9, 43 (2012).

[30] U. Brose, R. J. Williams, and N. D. Martinez, Allometric scaling enhances stability in complex food webs, Ecology letters 9, 1228 (2006).

[31] J. P. DeLong, B. Gilbert, J. B. Shurin, V. M. Savage, B. T. Barton, C. F. Clements, A. I. Dell, H. S. Greig, C. D. Harley, P. Kratina, et al., The body size dependence of trophic cascades, The American Naturalist 185, 354 (2015).

[32] K.-S. Cheng, S.-B. Hsu, and S.-S. Lin, Some results on global stability of a predator-prey system, Journal of Mathematical Biology 12, 115 (1982).

[33] S.-B. Hsu and J. Shi, Relaxation oscillation profile of limit cycle in predator-prey system, Discrete & Continuous Dynamical Systems-B 11, 893 (2009).

[34] C. Huffaker et al., Experimental studies on predation: dispersion factors and predator-prey oscillations, Hilgardia 27, 343 (1958).

[35] L. S. Luckinbill, Coexistence in laboratory populations of paramecium aurelia and its predator didinium nasutum, Ecology 54, 1320 (1973).

[36] F. K. Balagaddé, H. Song, J. Ozaki, C. H. Collins, M. Barnet, F. H. Arnold, S. R. Quake, and L. You, A synthetic escherichia coli predator–prey ecosystem, Molecular systems biology 4, 187 (2008).

[37] B. Blasius, L. Rudolf, G. Weithoff, U. Gaedke, and G. F. Fussmann, Long-term cyclic persistence in an experimental predator–prey system, Nature 577, 226 (2020).

[38] M. Trpis, Interaction between the predator toxorhynchites brevipalpis and its prey aedes aegypti, Bulletin of the World Health Organization 49, 359 (1973).

[39] S. Utida, Cyclic fluctuations of population density intrinsic to the host-parasite system, Ecology 38, 442 (1957).

[40] C. E. Goulden and L. L. Hornig, Population oscillations and energy reserves in planktonic cladocera and their consequences to competition, Proceedings of the National Academy of Sciences 77, 1716 (1980).

[41] R. Sudo, K. Kobayashi, and S. Aiba, Some experiments and analysis of a predator-prey model: Interaction between colpidium campylum and alcaligenes faecalis in continuous and mixed culture, Biotechnology and Bioengineering 17, 167 (1975).

[42] A. Rohatgi, Webplotdigitizer (2017).

[43] E. P. White, S. M. Ernest, A. J. Kerkhoff, and B. J. Enquist, Relationships between body size and abundance in ecology, Trends in ecology & evolution 22, 323 (2007).

[44] D. L. De Angelis, Estimates of predator-prey limit cycles, Bulletin of Mathematical Biology 37, 291 (1975).

[45] N. L. Lundström and G. Söderbacka, Estimates of size of cycle in a predator-prey system, Differential Equations and Dynamical Systems, 1 (2018).

[46] H. K. Khalil, Noninear Systems (Prentice-Hall, New Jersey, 1996).

[47] I. A. Hatton, K. S. McCann, J. M. Fryxell, T. J. Davies, M. Smerlak, A. R. Sinclair, and M. Loreau, The predator-prey power law: Biomass scaling across terrestrial and aquatic biomes, Science 349 (2015).

[48] U. Brose, T. Jonsson, E. L. Berlow, P. Warren, C. Banasek-Richter, L.-F. Bersier, J. L. Blanchard, T. Brey, S. R. Carpenter, M.-F. C. Blandenier, et al., Consumer–resource body-size relationships in natural food webs, Ecology 87, 2411 (2006).

[49] B. Gilbert, T. D. Tunney, K. S. McCann, J. P. DeLong, D. A. Vasseur, V. Savage, J. B. Shurin, A. I. Dell, B. T. Barton, C. D. G. Harley, H. M. Kharouba, P. Kratina, J. L. Blanchard, C. Clements, M. Winder, H. S. Greig, and M. I. O’Connor, A bioenergetic framework for the temperature dependence of trophic interactions, Ecology Letters 17, 902 (2014).

[50] J. R. Seymour, L. Seuront, and J. G. Mitchell, Microscale and small-scale temporal dynamics of a coastal planktonic microbial community, Marine Ecology Progress Series 300, 21 (2005).

[51] R. G. Eagon, Pseudomonas natriegens, a marine bacterium with a generation time of less than 10 minutes, Journal of bacteriology 83, 736 (1962).

[52] T. K. Lowenstein, B. A. Schubert, and M. N. Timofeeff, Microbial communities in fluid inclusions and long-term survival in halite, GSA Today 21, 4 (2011).

[53] T. J. Lavery, B. Roudnew, J. Seymour, J. G. Mitchell, V. Smetacek, and S. Nicol, Whales sustain fisheries: blue whales stimulate primary production in the southern ocean, Marine Mammal Science 30, 888 (2014).

